# How do backward walking ants (*Cataglyphis velox*) cope with navigational uncertainty?

**DOI:** 10.1101/2019.12.16.877704

**Authors:** Sebastian Schwarz, Leo Clement, Evripides Gkanias, Antoine Wystrach

## Abstract

Current opinion in insect navigation assumes that animals need to align with the goal direction to recognise familiar views and approach it. Yet, ants sometimes drag heavy food items backward to the nest and it is still unclear to what extent they rely on visual memories while doing so. In this study displacement experiments and alterations of the visual scenery reveal that ants do indeed recognise and use the learnt visual scenery to guide their path while walking backward. In addition, the results show that backward homing ants estimate their directional certainty by combining visual familiarity with other cues such as their path integrator and the time spent backward. A simple model that combines path integration with repulsive and attractive visual memories captures the results.

## INTRODUCTION

Central place foragers – such as desert ants – exhibit formidable navigational skills to find food and their way back home during numerous daily foraging trips (Collett, Graham, & Durier, 2003; Heinze, Narendra, & Cheung, 2018; Wehner, 2003). These ground dwellers rely on a set of navigational strategies such as path integration (PI) (Wehner & Srinivasan, 2003; Wittlinger, Wehner, & Wolf, 2006) and visual scene navigation (Cheng, Narendra, Sommer, & Wehner, 2009; Zeil, 2012). The literature agrees that ants continuously integrate the directional dictates of these different strategies together, rather than switching between them (Collett, 2012; Hoinville & Wehner, 2018; Legge, Wystrach, Spetch, & Cheng, 2014; Reid, Narendra, Hemmi, & Zeil, 2011; Wystrach, Mangan, & Webb, 2015).

Current models of insect visual navigation capture well the behaviour of forward navigating ants (Baddeley, Graham, Husbands, & Philippides, 2012; Hoinville & Wehner, 2018; Wystrach, Beugnon, & Cheng, 2011; Wystrach, Cheng, Sosa, & Beugnon, 2011; Zeil, 2012) however, how ants navigate while dragging a heavy food item backward remains unclear (Ardin, Mangan, & Webb, 2016; Pfeffer & Wittlinger, 2016; Schwarz, Mangan, Zeil, Webb, & Wystrach, 2017). Despite their irregular backward foot strides the ants’ PI system seems as accurate as during forward movement (Pfeffer, Wahl, & Wittlinger, 2016; Pfeffer & Wittlinger, 2016), however, guidance based on terrestrial visual cues seems disrupted (Schwarz et al., 2017). Evidence suggests that to recognise the familiar terrestrial scenery ants need to align their body in the familiar forward direction (Narendra, Gourmaud, & Zeil, 2013; Wystrach, Cheng, et al., 2011; Zeil, 2012). This is probably why ants dragging a food item backward occasionally display a so-called ‘peeking’ behaviour: the ant stops pulling, drops its food item and turns around to look forward. If the scenery is familiar, the ant quickly returns to her food item and adjusts her backward path in the newly corrected homing direction. It seems clear that during these few moments facing forward in a familiar direction, ants recover and store the correct direction; and subsequently rely on celestial cues to maintain this new bearing when traveling backward (Schwarz et al., 2017). In this case navigation is discretised into different sources of information being used sequentially rather than simultaneously. Also, ‘peeking’ involves the decision to trigger a distinct and observable behaviour when navigational information is needed. This behaviour therefore provides a good opportunity to investigate how ants estimate their navigational uncertainty and as a corollary, which navigational information they have access to.

Here two experiments with backward walking ants were carried out to investigate the following questions: (1) Can ants still perceive visual familiarity when walking backward? (2) How can this visual information enable ants to control their backward path. (3) Which information is used by ants to estimate uncertainty and trigger a peeking behaviour?

## METHODS

### Study animal and site

The experiments were carried out with Spanish desert ants *Cataglyphis velox* on a field site with diverse grass and bush vegetation at the outskirts of Seville during June 2017 and 18. *Cataglyphis velox* show typical characteristics of a desert ant such as diurnality, thermophily and solitary foraging (Cerda, 2001). As in other ant species, navigation and orientation in *C. velox* is predominantly based on vision derived from terrestrial and celestial cues (Mangan & Webb, 2012; Wystrach et al., 2015).

### General methods

Two experiments were conducted: Experiment 1 in 2017 and Experiment 2 in 2018. Both set-ups shared the following methods.

Ants were restricted to forage on a straight route between their nests and a feeder. The routes were mostly cleared from vegetation and enclosed by thin white plastic planks (10 cm high) that were dug halfway into the ground. The slippery surface of the planks prevented ants foraging elsewhere while minimising the obstruction of surrounding views (Wystrach, Beugnon, & Cheng, 2012). Ants could freely travel between nest and feeder, which was a ~15×15×15 cm plastic bowl sunk into the ground that contained several kinds of sweet buttery biscuit crumbs. The walls of the bowl were covered with a thin layer of Fluon® and prevented ants from climbing. Ants that dropped into the feeder and picked up a crumb were marked individually with coloured acrylic or enamel paint (Tamiya™). During training, ants could leave the feeder via a small wooden ramp. Ants were considered trained and ready for testing once they had performed at least five foraging runs and were able to reach the feeder from the nest in a straight line (without colliding into any barriers).

During tests (see below) the feeder ramp was removed to prevent other homing ants from interfering.

### Experiment 1

Experiment 1, conducted during summer 2017, entailed a nest at the beginning of a straight 8×1.8 m long foraging route. Three large wooden boards (2.4×1.2 m) were connected (7.2×1.2 m) and placed onto the foraging route. These boards enhanced the tarsi grip of the ants and provided an even substrate that minimised potential interference with small grass haulms or pebbles during tests when ants dragged their food items backward (Fig. 1a).

**Figure 1.**
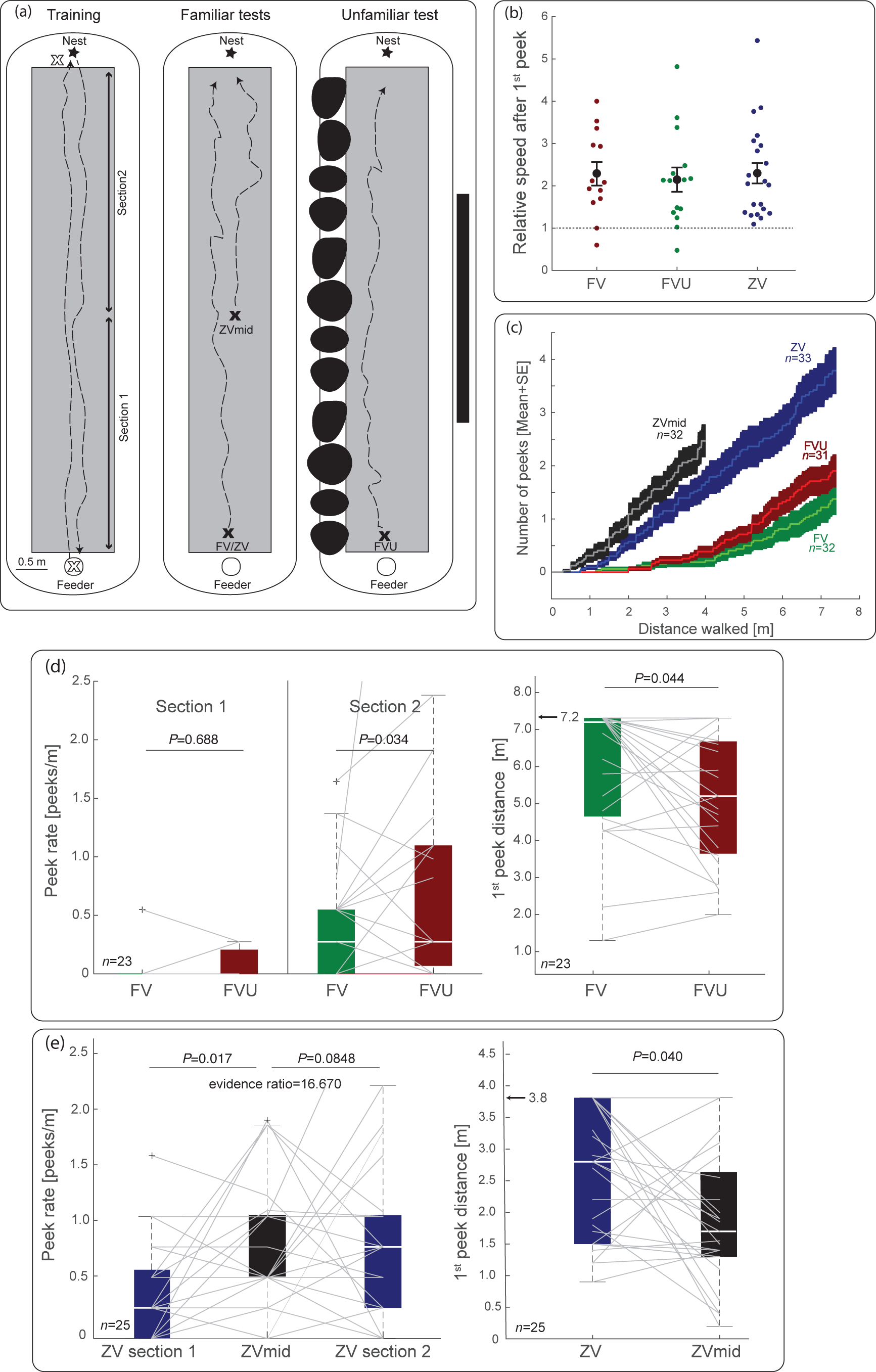
Dynamics of peeking behaviours in terms of path integration and visual familiarity (Experiment 1). (a) Schematics of experimental set-up with training and test conditions. During training ants foraged between nest and feeder (~8.0 m) on three thin wooden boards serving as an even substrate (grey rectangles). The route was divided into two sections corresponding to the first and second half of the route. For tests, trained ants were either captured at the feeder (full-vector ants, FV) or just before entering the nest after foraging (zero-vector ants, ZV; open crosses) and released at the feeder (as FV-FVU or ZV-ants) or on the middle of the route (ZVmid-ants; black crosses). For FVU (unfamiliar), the familiarity of the route was manipulated by adding large black visual objects (black blobs) on one side and a dark tarp (black vertical bar) on the other side of the route. Dashed lines depict example paths of the ants. (b) Change in speed after the first peek. Each dot shows the relative change in speed (5s after/5s before) thefirst peek for each ant. Dotted line at 1 indicates no change in speed. Almost all ants increase their speed after their first peek. (c) Cumulated number of peeks displayed against the distance walked along the route (M±SE across individuals). A clear separation between FV-and ZV-ants is visible where ZV-ants peek earlier than FV-ants. (d) Overall peek rate (left) and distance of the first peek (right) for both FV-ant conditions. FV-ants peek less often than FVU-ants in Section 2 and travel a longer distance before displaying their first peek. 7.3 m indicates the end of the route. (e) Overall peek rate (left) and distance of the first peek (right) for both ZV-ant conditions. ZVmid-ants peek more often than ZV-ants from Section 1 but not from Section 2 (Bayesian evidence ratio strongly favour similarity with Section 2). ZVmid-ants travel a shorter distance before first peek as compared to ZV-ants. 3.8 m indicates the end of the route (ZV-ant paths were truncated at 3.8 m to match the maximum homing distance of ZVmid-ants). Grey lines (d, e) represent individually tested ants across conditions. See main text for statistical details.

During training, the individually marked foragers scuttled (forward) between the nest and feeder over the connected boards and familiarised themselves with the visual surroundings. After training, individual ants were subjected to one test conditions. All tests comprised of a forager that dragged a large biscuit crumb backward. For that, trained foragers with a small food item (~0.2×0.2×0.2 cm) were caught and transferred into a plastic vial. The food item was carefully and manually removed and a larger biscuit piece (~2.0×0.5×0.2 cm) was offered to the ant instead. The biscuit provided was large enough to force the ants to drag it backward. After the ant locked mandibles onto the large biscuit, she was transferred to the appropriate release point. Four possible test conditions were carried out with either FV-(i.e., ants with their PI vector information, captured at the feeder) or ZV-ants (i.e., foragers without PI information captured just before they enter the nest; Fig. 1a). To test the effect of the level of visual familiarity in backward movements, ants were either released at the familiar feeder (FV) or at the feeder with unfamiliar visual surroundings (FVU). A few seconds after the FVU-ant had started to home backward the visual surroundings were altered by adding large black plastic bags (~0.8×0.6 m) on one side and a large dark tarp (0.9×3.4 m) on the other side of the route. The objects were always placed parallel to the backward path of the ants to avoid behavioural interferences and potential obstructions. To test the effect of route location, backward moving ants were tested either at the feeder (beginning of the route; ZV) or at the middle of the route (ZVmid).

### Experiment 1: data and analysis

For all tests, the distance between the release point and the location at which peeking behaviours occurred was noted. Tests ended as soon as the backward walking ant reached the end of the wooden boards (i.e., ~0.5 m in front of the nest entrance) or abandoned her food item for more than one minute. Individual ants were tested only once per test but were subjected to different test conditions with at least one un-interfered training trial between tests. The sequence of tests was evenly counter-balanced across individuals.

Comparison were made between FV-vs. FVU-ants and ZV-vs. ZVmid-ants (Fig. 1a). Given the large inter-individual variations, paired-data was applied and thus only ants that were tested on both FV and FVU or ZV and ZVmid conditions, respectively were kept for analysis. Both the distance at which the first peeking behaviour occurred (1^st^ peek distance) and the overall peek rate of individuals (i.e., number of peek/distance walked) were compared using Wilcoxson ranksum tests a nonparametric statistic for paired data (Matlab™, Mathworks, Matick, MA, USA).

Given that all ants walked rather straight toward the nest along the route, distance walked could be simply approximated by the beeline distance walked along the route. Most ants walked the full route (7.2 m) except obviously ants in the ZVmid condition and some foragers that abandoned their biscuit. For the comparison of peek rate, the 7.2 m long route was divided approx. into half (Section 1: 0 - 3.4 m; Section 2: 3.4 - 7.2 m; Figure 1A). Thus, during ZVmid tests ants ran only Section 2. Comparisons between ZV-vs. ZVmid-ants were conducted to separate the effect of distance walked (i.e., ZVmid vs. ZV on Section 1) from the actual location along the route (i.e., ZVmid vs. ZV on Section 2). Bayesian statistics were applied to evaluate which of these alternative hypotheses explain peek rate best.

Backward paths were recorded by using GoPro HERO3+ cameras which were manually held approx. 0.6 m above the tested ant. Therefore, a quantification of the movement speed of the ants before and after peeking could be calculated. For that the relative distance walked by the backward ants during the five seconds preceding the onset of the first peek (i.e., before the moment when the ants released the biscuit) and five seconds after the peek (i.e., after the ant resumed backward motion) was estimated.

### Experiment 2

Experiment 2 was conducted in the summer months of 2018 with two different nests of *C. velox* ants. For each nest, a 5.0×2.0 m straight foraging route was built with the nest entrance at one end and the feeder at the other end (Fig. 2a). As in Experiment 1, the route was enclosed by white plastic planks and ants were given a choice of biscuit crumbs inside the feeder to prompt foraging. However, here the ants scuttled back and forth directly on the natural ground during training. Once trained (see General methods), individual ants were captured on their way home 0.5 m before reaching their nest and subsequently released at one out of four possible locations (Fig. 2a):

**Figure 2.**
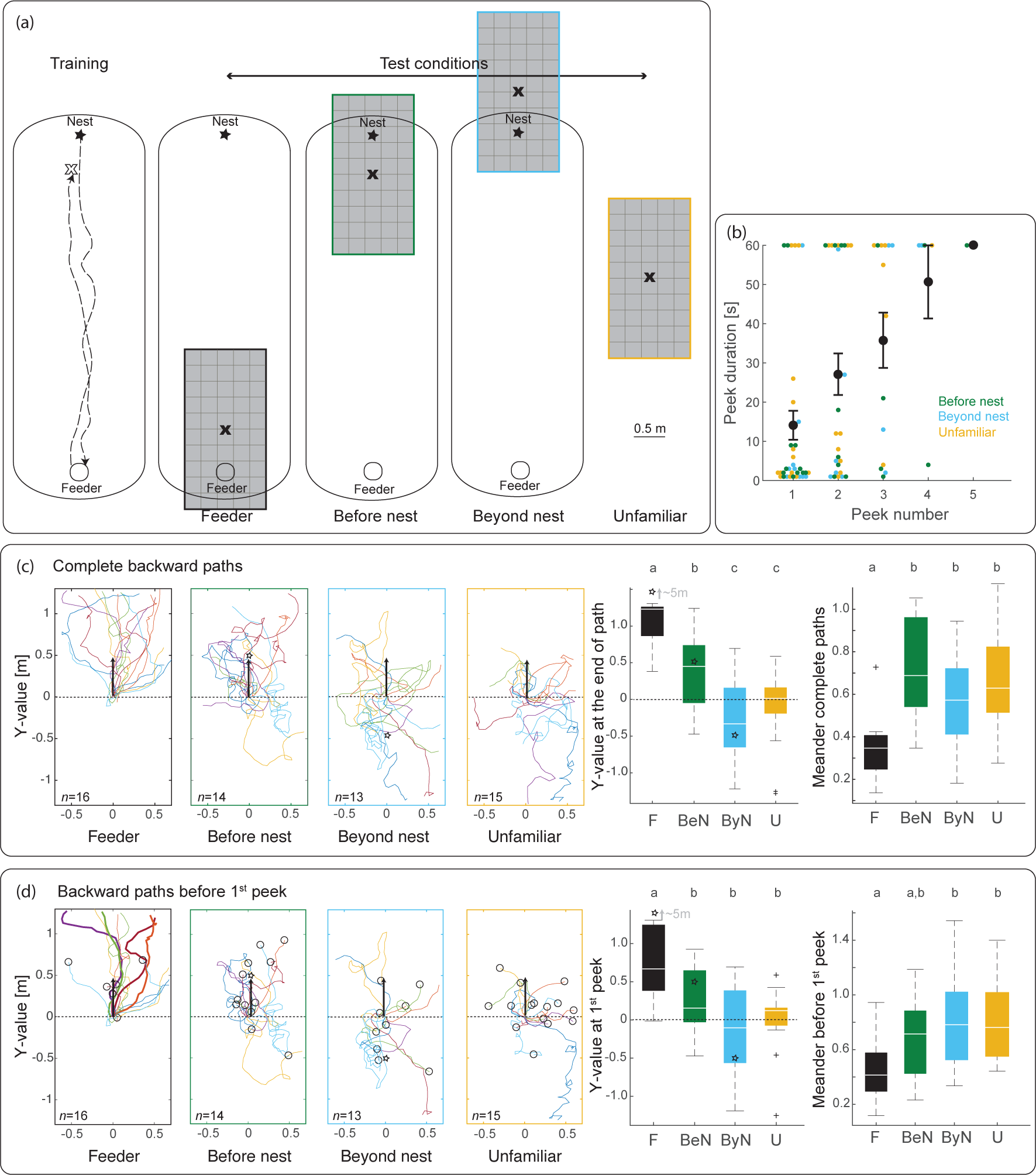
Backward path characteristics and peeking behaviour at different release points (Experiment 2). (a) Schematics of experimental set-up with training and test conditions. During training ants foraged between nest and feeder (~5.0 m). Trained ants with a remaining homing vector of 0.5 m (open cross depicts capture point) were tested backward at different release points: feeder (F), before nest (BeN), beyond nest (ByN) and unfamiliar (U) test site. Ants were released (black crosses) on the middle of a thin wooden board (grey) to rule out the use of olfactory cues. Boards had a 0.25×0.25 m squared pattern to enable path recording. Dashed lines depict example paths of ants during training. (b) Duration of peek (the time the ant spent away from the cookie) as a function of peek number for each individual. Regardless of test condition, peek duration increases with number of peeks. Max. threshold of peek duration was set to 60s and ‘Feeder condition’ was removed from this analysis as peek number correlates with position along the board. (c) Complete recorded backward paths of ants across conditions. Paths ended either because ants left the board or abandoned their cookie (peek duration > 60s). Boxplots show distance reached at the end of the path along the feeder nest axis (Y axis) and meander of the paths across individuals. Differences in top letters (a, b, c) indicate significant differences between groups (alpha=0.05). Except for unfamiliar (U) all other conditions were directed toward the actual nest position, showing that ants used familiar visual cues. Black dotted lines depict release point, black arrows remaining vector length (0.5 m) and open stars actual nest position. (d) As in (c), except that paths were truncated at the first peek or when exiting the board. Hence all navigational information gathered by ants was obtained while walking backward. Thick paths (left panel) emphasise four ants in the feeder conditions that displayed nest-directed backward paths without facing the nest direction. Open circles represent positions of 1^st^ peeks. For statistical details see text.

Feeder (F): Ants were released 0.5 m after the feeder.

Before nest (BeN): Ants were released on the route, 0.5 m before their nest. Beyond nest (ByN): Ants were released 0.5 m beyond the nest in the feeder-nest direction.

Unfamiliar (U): Ants were released ~30.0 m away from the nest in a visually unfamiliar location.

For all tests, ants were captured in a plastic vial, offered a large biscuit crumb to incite backward walking and, once the ant had grabbed the cookie, released within a lampshade at the middle of a large wooden board (2.4×1.2 m). Individual ants were tested only once per test, but could be subjected to different tests conditions, with at least one un-interfered training trial between tests. The wooden board was set in place just before and removed just after each test. The board was centred on the current release location with the long side along the feeder-nest direction (Fig. 2a) as it represents the expected homing direction. The board provided an even substrate during tests and prevented ants to use potential familiar olfactory cues from the ground or the nest (BeN and ByN tests). A grid pattern (0.25×0.25 m) drawn on the board enabled paths to be transcribed onto gridded paper. The lampshade (beige opaque fabric, 0.5 m diameter; 0.4 m height) surrounded the ant upon release and obstructed any familiar terrestrial views; the top of the lampshade was open providing the ant with celestial compass cues. Once the ant had pulled the large crumb backward for 0.1 m, the lampshade was lifted and the visual surrounding was revealed. This ensured that the ants could not utilise any familiar scenes before starting their backward path.

The backward paths and locations of peeking behaviour were noted. For each peek, the duration (i.e., the time the ant was not dragging the biscuit) was recorded but not the forward paths during the peek itself. Recording continued until the ants either reached the edge of the board or abandoned their large crumb for more than 1 min.

### Experiment 2: data and analysis

The recorded paths were digitised as (x, y) coordinates using GraphClick (Arizona Software). Path characteristics such as direction, meander (for details see Schwarz, Albert, Wystrach, & Cheng, 2011) and peek location were computed and analysed with Matlab™ (Mathworks, Matick, MA, USA). Differences between test locations were determined using a generalised linear mixed effect model with repeated ants as random effect and Tukey’s post hoc corrections. For peek durations, a model for proportional (binomial) data was applied with 0 to 60 s (the duration at which we stopped recording) reported between 0 and 1.

## RESULTS

### Experiment 1

In this experiment ants were free to scuttle forward along an 8.0 m straight route between the nest and a feeder to become familiarised with the visual scenery of the route. For tests, trained homing ants were captured either directly at the feeder (FV-ants) or upon reaching their nest (ZV-ants). Captured ants were given a large biscuit crumb that had to be dragged backward along the route home (Fig. 1a) and the occurrence of peeking behaviour was recorded.

#### FV vs. ZV: effect of path integration

In both FV and ZV conditions, ants started to pull their biscuit toward the nest and mostly maintained that direction. The ability of ZV-ants to do so suggests that the foragers were able to perceive the familiar terrestrial cues although they might have had a chance to take a glimpse forward upon release and before starting their backward movements. In any case, the PI state had a strong effect. As seen by the standard errors of the data population, ZV-ants peeked earlier (first peek distance ZV-ants: M±SD = 3.28±2.19 m; FV-ants: M±SD = 5.90±1.93 m) and thrice as much (overall peek rate ZV-ants: M±SD = 0.63±0.63 peek/m; FV-ants: M±SD = 0.19±0.29 peek/m) than FV-ants (Fig. 1c). Also, ZV-ants occasionally abandoned their large food item and did not resume backward movements (6 out of 33), whereas no FV-ants abandoned theirs (0 out of 32). A Fisher’s exact test verified a significant difference (*P* = 0.032). It seems clear that a lack of (or conflicting) PI information decreases the ants’ directional certainty.

#### FV vs. FVU: effect of visual unfamiliarity

To test the potential effect of the level of visual unfamiliarity on backward walking ants, two conditions were conducted: (1) FV-ants homing backward on the unaltered, usual route, and (2) Full-Vector-Unfamiliar (FVU) ants, homing backward on the same route but this time the visual surrounding was altered by additional large black plastic bags (~0.8×0.6 m) and a rectangular dark tarp (0.9×3.4 m) on each side of the route (Fig. 1a). The objects were added only after the tested FVU-ants had started their backward path to ensure that they could not monitor the visual change before engaging in dragging the biscuit. If ants trigger peeks because of navigational uncertainty then they should peek more often in unfamiliar environments. Results confirm the prediction.

First, FVU-ants peeked more often than FV-ants. However, this effect was weak, and reaches significance only in Section 2 (Wilcoxson ranksum test: *P* = 0.027, *Z* = 3.751) but not in Section 1 of the route (Wilcoxson ranksum test: *P* = 0.688, *Z*~0; Fig. 1d), due to a statistical floor effect. Indeed, a low rate of peeking in the first section of the route was expected, given that the path integration vector is longer and thus stronger at the beginning of the route home (Wystrach et al., 2015).

Second and most importantly, FVU-ants travelling in the unfamiliar environment displayed their first peek earlier along the route as compared to FV-ants on the familiar route (Wilcoxson ranksum test: *P* = 0.044, *Z* = 2.016; Fig. 1d). The results suggest that ants could perceive the difference in visual familiarity while walking backward given that the visual surrounding was altered only after the ants had started they journey backward,

As for ZV-ants (see above), FVU-ants tested in the unfamiliar condition abandoned their biscuits significantly more than FV-ants (FVU: 6 out of 31 vs. FV: 0 out of 32. Fisher’s exact test: *P* = 0.022). Here again, it seems that visual unfamiliarity decreases directional certainty of backward walking ants.

#### ZV vs ZVmid: effect of location

We investigated the potential effect of the location along the route by releasing zero vector ants either at the beginning of the familiar route (ZV) or directly in the middle of the familiar route (ZVmid; Fig. 1a, c). Consequently, ZVmid-ants walked only Section 2, while ZV-ants moved along both sections. Ants displayed their first peek on average slightly earlier when released at the middle of the route (ZVmid) than when released at the beginning of the route (ZV; Wilcoxson ranksum test: *P* = 0.040, *Z* = 2.062; Fig. 1e). Also, the peek rate displayed by ZVmid-ants along Section 2 (the only section they walked) was higher than ZV-ants along Section 1 (Wilcoxson ranksum test: *P* = 0.005, *Z* = −2.814) but similar to the peek rate displayed by these ZV-ants along Section 2 of the route (Wilcoxson ranksum test: *P* = 0.796, *Z* = - 0.2585; Fig. 1e). A Bayesian evidence ratio was computed to estimate whether Section 1 or Section 2 of ZV-ants’ peek rate resembles most ZVmid-ants’ peek rate. The obtained evidence ratio was 50.74 in favour of Section 2, which equals ‘overwhelming evidence’ for an effect on peek rate of the actual location along the route rather than the distance walked.

#### Peeking and walking speed

Interestingly, in all conditions, and for the vast majority of the individuals, ants walked backward on average twice as quickly after peeking than before peeking (Fig. 1b).

This supports the idea that a peeking event increases the ant’s directional certainty for some time.

### Experiment 2

In this experiment, homing foragers were trained along a route, captured 0.5 m before they reached their nest, provided with a large biscuit crumb and released on top of a wooden board (Fig. 2a) at different test locations: namely, 0.5 m after the feeder (F), 0.5 m before the nest (BeN), 0.5 m beyond the nest (ByN) and at a distant unfamiliar location (U) ~30.0 m away (Fig. 2a). Crucially, in this experiment all tested ants were prevented from monitoring the visual surrounding before dragging their food item backward as a lampshade was blocking the whole panoramic view (see Methods). Hence, any evident effect of the scenery on the backward path must result from visual information perceived while the ants were dragging their crumb backward – at least until they peeked for the first time.

#### Ants can guide their backwards path

Analysis of the complete paths trajectory revealed differences between test conditions. The Y-values – the position along the feeder-to-nest line – at the end of the foragers’ path varied across conditions (ANOVA: *F*=21.96, *P*<0.001; Fig. 2c). Ants from the feeder test displayed paths directed toward the nest and hence obtained higher Y-values at the end of their recorded paths than any other conditions (Tukey’s post-hoc test F vs. BeN, ByN and U: *Z*s > 3.75, *P*s < 0.001; Fig. 2c). Ants in unfamiliar tests showed no directional preference along the Y-axis (Fig. 2c), as expected given the lack of familiar visual information at this location. Interestingly, ants from BeN and ByN conditions differed significantly in their final Y-values on the board (Tukey’s post-hoc test BeN vs. ByN: *Z* = 3.47, *P* = 0.003). The medians of both of these groups are close to the actual nest location, showing that they used familiar visual cues to search at the nest (Fig. 2c). Differences between conditions could also be observed in path meander (ANOVA: *F*=9.07 *P* < 0.001). Ants from the feeder test showed straighter paths than ants from all other conditions (Tukey’s post-hoc test F vs. BeN, ByN and U: *Z*s > 3.68 *P*s < 0.002; Fig. 2c). No difference in meander among the remaining test conditions could be determined. Indeed, BeN, ByN and U ants were expected to search on the board: BeN-and ByN-ants due to the proximity of the nest and U-ants due to the of the lack of familiar visual information. Overall, these data show that ants could use familiar visual cues to adequately direct their backward paths.

Remarkably, analysis of the paths displayed before the first peek (or until the ant left the board if she did not peek) showed a similar pattern of results for both distance reached along the Y-axis (ANOVA: *F* = 11.37 *P* < 0.001) and path meander (ANOVA: *F* = 3.52 *P* = 0.024). Ants released at the feeder travelled significantly longer distances along the feeder-nest direction before peeking than all other test conditions (Tukey’s post-hoc test: F vs. BeN, ByN and U: *Z*s > 3.29 *P*s < 0.006; Fig. 2d) and displayed straighter paths (Tukey’s post-hoc F vs. ByN and U: *Z*s > 2.65 *P*s < 0.03; F vs. BeN: *Z* = 1.88 *P* = 0.235; Fig. 2d). The three other groups (BeN, ByN, U) were expected to search on the board and to perform a similar level of path meander.

Differences in the feeder-nest distance between these conditions (BeN, ByN, U) were not significant using Tukey’s post-hoc test. However, the pattern of results followed what was expected if ants were using views to direct their path toward the nest. Ants released before (BeN) and beyond (ByN) their nest both moved on average toward the nest location, that is, in opposite direction from their release points; and ants released at the unfamiliar test site (U) showed less directed paths (Fig. 2d). The differences in paths characteristics is also reflected if one considers the probability of obtaining the expected order of path endpoint across the four test conditions (Y-value: F > BeN > U > ByN) is 1/4! = 0.042. Interestingly, several ants released at the feeder (4 out of 16 ants) displayed nest-directed backward paths across the whole recording board without performing a single peek and by keeping their body orientation away from the feeder-nest direction by at least 90° (bold paths in Fig. 2d).

Because nest-directed path sections were achieved before the ants triggered their first peek and the visual panorama was revealed to them only after they had started backward motion, the differences across locations show that ants can recognise and use the familiar visual cues to guide their path while moving backward and without the need of peeking.

#### Peek duration and past information

We also tested whether peek duration was influence by the test condition and the number of previously displayed peeks (Fig. 2b). The feeder condition was excluded from this analysis as these ants were expected to move in a straight line and exit the board so that the actual peek number of a given ant may correlate with the location where the ant peeks instead of being based on the previous peek(s): the higher the peek number the larger the distance from the feeder. The three other groups (BeN, ByN, U) on the other hand, are expected to search on the board so any effect of the peek number is unlikely to be attributed to a specific location on the board.

Interestingly, peek duration, which was recorded up to 60 s, was strongly influenced by the number of peeks previously displayed by the ant (GLM peek number: *F* = 17.09, *P* < 0.001; Fig. 2b) and not the actual test condition (GLM condition: *F* = 0.17, *P* = 0.841; Fig. 2b). The more peeks an ant had previously displayed the longer its current peeking duration. This shows that the ant’s peeking behaviour is modulated by past information but whether it is the time passed or the number of peek previously displayed cannot be disentangled here.

## DISCUSSION

Ants dragging a heavy food item backward occasionally trigger a so-called ‘peeking behaviour’ or ‘peek’: ants drop their food and turn around to look forward. Aligning their body in a familiar direction enables them to recognise the learnt visual panorama and hence adjust the direction of their subsequent backward path (Schwarz et al., 2017). It is clear that ants gain directional information from learnt terrestrial cues when peeking forward. However, whether or not they can recognise terrestrial cues while dragging their food item backward is less clear. Several of the current results demonstrate that ants are indeed able to do so, raising question about the underlying mechanisms.

### Ants still recognise terrestrial cues while walking backwards

Experiment 1 shows that the visual scenery experienced while walking backwards influenced the occurrence of peeking behaviour. First, ZVmid-ants displayed their first peek earlier when starting their backward journey halfway along the route rather than ZV-ants at the beginning of the route (Fig. 1e). Second, FV-ants displayed their first peek earlier along the route if the surrounding scenery was artificially altered (FVU, Fig. 1d). This was true even though the scene was manipulated only after the ants had started dragging their biscuit backward and thus indicates that ants perceived the alteration of the familiar scene while walking backward. It should be noted that this effect was weak (Fig. 1d), possibly because the alteration of the scene was not obvious enough (Schwarz et al., 2014).

In Experiment 2, ants could guide their trajectories based on terrestrial cues while walking backward. Ants were released on a board (ruling out the use of olfactory cues) and within a lampshade. The visual world was revealed to them once they had started their backward journey. Nonetheless and despite the lack of PI homing vector, their paths were oriented in the expected direction (i.e., the nest) resulting in differences between test conditions. Importantly, this was also true for the portion of path displayed before their first peek, that is, displayed purely backward (Fig. 2d).

In sum, ants can use learnt terrestrial visual cues while walking backward to guide their path as well as decide whether and when to peek forward. The next section discusses potential explanations.

### Mental rotation or combining attractive and repulsive views?

How can ants recognise views backward? This is a puzzling question given that the assumption of current models of visual homing states that views must be retinotopically aligned to provide directional information (Ardin, Peng, Mangan, Lagogiannis, & Webb, 2016; Baddeley et al., 2012; Collett, Graham, & Collett, 2017; Möller, 2012; Wystrach, Mangan, Philippides, & Graham, 2013; Zeil, 2003). This idea seems to be supported by data in freely navigating ants (Narendra et al., 2013; Wystrach, Cheng, et al., 2011) although some other processes may be also at work (Wystrach et al., 2012). Recently, it has been suggested that ants may perform some sort of mental rotation to compare misaligned views (Ardin, Mangan, Wystrach, & Webb, 2015; Ardin et al., 2016), which could be achieved if views are encoded in the frequency domain (Stone et al., 2017). But, this idea is hard to reconcile with the result of previous experiments where ants would not adjust their backward trajectory at all unless they peeked to align their body in the correct direction (Schwarz et al., 2017).

Here we suggest an alternative hypothesis to mental rotation: ants may still need to align their body to recognise views retinotopically but possess a memory bank of views learnt while facing in multiple directions and not only toward the nest. Notably, views learnt while facing in the anti-nest direction could be treated as repulsive when homing (Fig. 3a). The familiarities resulting from the comparison of the currently perceived view with both attractive (nest facing) and aversive (feeder facing) visual memories could simply be compared in a way somewhat analogous to an opponent process. The signal resulting from this comparison informs the ant about whether to move toward or away from the currently faced direction. In addition, homing ants might use the visual memories stored during their outbound trips (i.e., when they went from the nest to the feeder) as repulsive. This idea challenges the opinion that ants treat in-and outbound trip visually separately depending on the motivational state (Harris, Hempel de Ibarra, Graham, & Collett, 2005; Wehner, Boyer, Loertscher, Sommer, & Menzi, 2006). Instead, ants may always recall both their memorised in-and outbound facing views but treat them as repulsive or attractive depending on their current motivational state.

**Figure 3.**
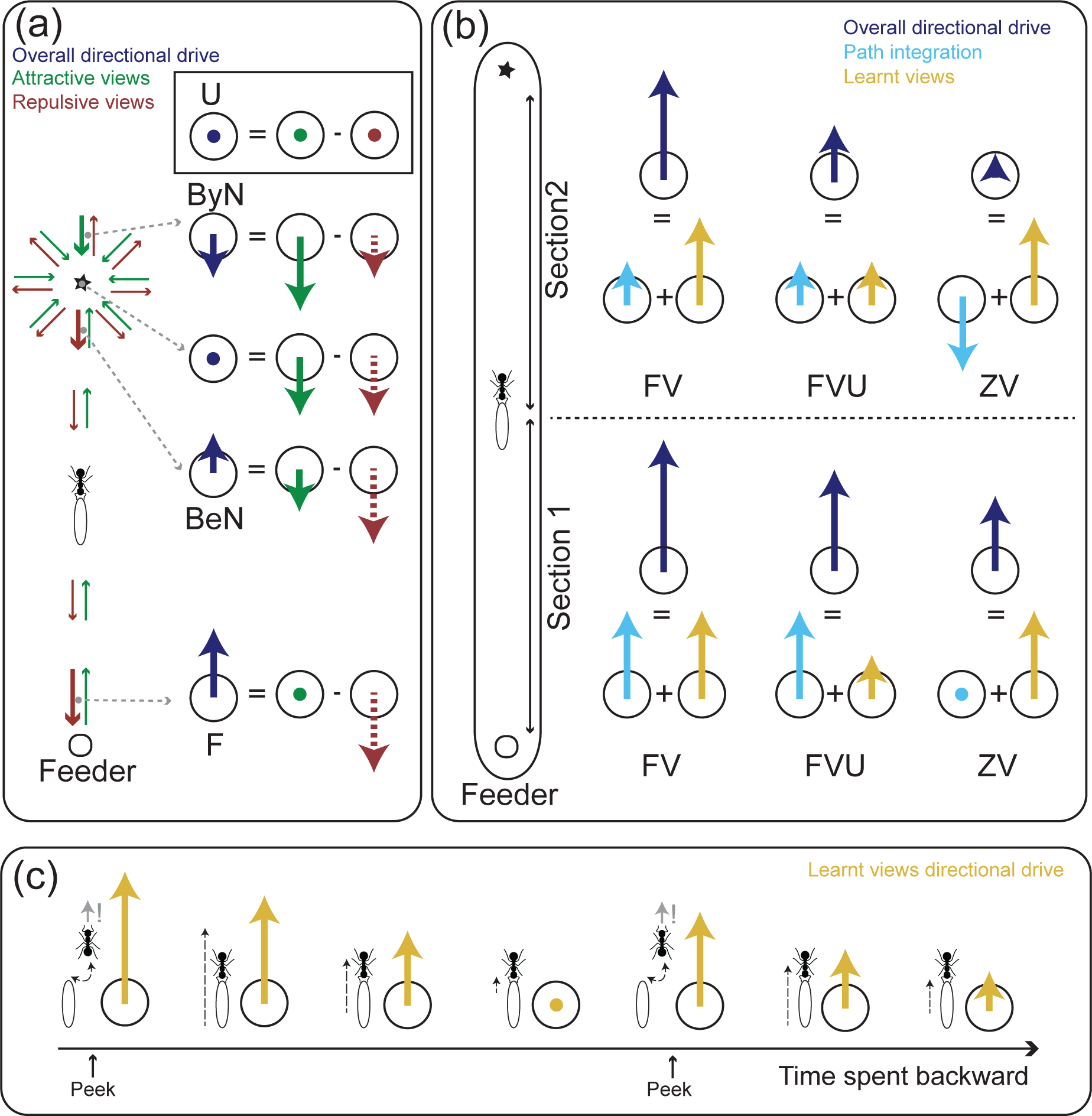
Models on how ants may combine navigational information. The longer the length of the drawn vectors the higher the directional drive. Dots represent cases with no directional drive and stars nest locations. (a) Illustration of the ‘repulsive view hypothesis’. Overall directional drive results from the integration of attractive homing views (green arrows) minus repulsive outbound views (red arrows). Left scheme represents theoretical positions and orientations of memorised views. The ant recognises only views that are aligned with its current body orientation (here, always facing downward). The larger the distance from the current location to the closest aligned view, the lower the familiarity and directional drive. This principle provides appropriate guidance toward the nest. Right scheme shows examples of integration for different locations (grey dashed arrows) with the ant always facing downward. For instance, when facing downward at the BeN location, the closest aligned view is repulsive (bold red arrow on right left scheme). Given that the neighbouring green arrow is not aligned, the closest aligned attractive view is further away, beyond the nest (bold green arrow on left scheme). Overall, the ants at this position (still facing downward) will be more repulsed than attracted by the current facing direction and thus walk backward toward the nest. In contrast, when beyond the nest ByN (still facing downward), the attractive views will match better than the repulsive views and the ants will thus turn around and walk backward toward the nest. Whatever the position and orientation of the ants around the nest, the agent will be drawn towards the nest (b) Directional drive across test conditions of Experiment 1 in Section 1 and 2. Ants are tested in FV (full-vector), FVU (full-vector unfamiliar) and ZV (zero-vector) conditions. Overall directional drive (dark blue vectors) results from the integration of path integration (cyan vectors, the longer the path integration vector the stronger its directional drive) and learnt view (yellow vectors, the more familiar the view the stronger its directional drive). (c) Directional drive resulting from the recognition of a learnt view (yellow vectors) decrease with time spent facing in a different direction. Low directional drive results in lower speed (dashed arrow) and eventually peeking. Here memorised views are assumed to be facing upward and are thus recognised only when facing upward during peeking (small grey arrow) and not while facing downward during backward motion. Note that the second peek triggers a lower directional drive than the first (see also Fig. 1b).

Such a hypothesis explains several observed phenomena of the current study. (1) In a former experiment (Schwarz et al., 2017) backward ants were not able to correct their path at all while walking backward because, in this particular set-up, in-and out-bound routes were spatially separated (as a one-way circuit) so that no outbound views where available to potentially help out backward homing ants. (2) In the current Experiment 2, backward ants released at the feeder (F) carried on in the correct nest direction (Fig. 2d) because they recognised outbound views oriented toward the feeder, driving them away from (or opposite to) this direction (Fig. 3a). (3) Alteration of the visual surrounding would trigger earlier peeking behaviours because the familiarity of the feeder facing (outbound) views would be equally altered, disrupting the repulsive effect and thus reducing the overall directional drive (Fig. 1d). (4) Assuming that outbound views near the feeder are more familiar than in the middle of the route (ants perform learning walks at the feeder: Nicholson, Judd, Cartwright, & Collett, 1999), the repulsive effect would be stronger for ants released at the feeder than in the middle of the route, yielding to a stronger directional drive and hence less peeking near the feeder (Fig. 1d). (5) Further, it was surprising in Experiment 2 that ants released close to the nest could direct their backward paths toward the nest (BeN, ByN; Figure 2c, d). Although this verifies that they recognised familiar views they nonetheless tended to peek often and even abandoned their cookie close to the nest (9 out of 28 in BeN-and 4 out of 20 in ByN-test). This seems counter-intuitive, yet it can be explained in the light of the ‘repulsive view’ hypothesis. During learning walks around the nest, ants appear to store indeed both nest and anti-nest oriented views (Jayatilaka, Murray, Narendra, & Zeil, 2018). Even if these nest views may be all extremely familiar, the integration of attractive (nest-oriented) and repulsive (anti-nest oriented) views would result in a low overall directional drive, which would thus lead to high peek rates (and a high probability for abandoning the crumb) but nonetheless guide the ants toward the nest. Fig. 3a illustrates further the functionality of this simple model. (6) Finally, two recent studies revealed the importance of outbound trips for homing ants. Ants with outbound views during training were more efficient in homing than ants with only inbound views during training (Freas & Cheng, 2018; Freas & Spetch, 2019). But whether homing ants actually used their outbound view as suggested here, or simply learnt homing views by turning back during their outbound trips remains to be seen.

### Ants combine multiple cues to estimate directional uncertainty and trigger peeks

It is known that ants combine the directional dictates of the current visual familiarity with their PI in a weighted fashion (Wehner, Hoinville, Cruse, & Cheng, 2018). Notably, the direction indicated by the current view is more weighted as the current view is familiar (Legge et al., 2014) and the direction indicated by the PI is more weighted as the PI vector length increases (Wystrach et al., 2015). Backward walking ants appear to weight these cues in the same fashion. Fig. 3b shows how such an integration of cues captures the peek rate observed across our conditions and distance walked along the route. Notably, this explains why ZV-ants peeked earlier and more often than FV-ants (Fig. 1b), as observed in North African *Cataglyphis fortis* ants (Pfeffer & Wittlinger, 2016) and why peek rate increases as the distance along the route increase (Fig. 1d, e).

Interestingly, such an estimate of directional certainty seems not only to influence the occurrence of peeking but also whether the peeking ants decided to return to their biscuit or abandon it. FV-ants in familiar visual surroundings and therefore with the highest directional certainty, dragged their biscuit all along the 7.20 m route without exception (32 out of 32). In contrast, some ants abandoned their biscuits in both FV unfamiliar (6 out of 31) and ZV (6 out of 33) conditions.

Finally, it is worth mentioning that ants clearly increased the speed of their backward motion after peeking (Fig. 1c). The increase in speed is likely not only a mere consequence of a short recovery from the dragging activity but also a gain in navigational certainty as this sudden speed increase is also apparent when displaced foragers reach their familiar route and recognise the familiar scenery (pers. observ.). Furthermore, a decrease in speed when ants run off their PI has been observed (Buehlmann, Fernandes, & Graham, 2018). Hence it seems that the speed of movement reflects the strength of the directional drive and therefore directional certainty. Taken together, peeking behaviour seem to increase directional certainty, at least temporally (Fig. 3c)

### Ants gather information about the time spent backward

Recently, it has been shown that directional information based on terrestrial cues is obtained when ants face forward during peeks and must therefore be stored in a short-term memory while the ant is subsequently walking backward (Schwarz et al., 2017). Our results suggest that short-term memory also influences the ants’ navigational certainty. The time spent peeking forward varied (as already noted by Pfeffer and Wittlinger (2016)) with some ants ‘peeking’ for less than a second while other spent more than 60s (after which recording was stopped) without returning to their cookie. Interestingly, the more an ant had peeked before during a test run the longer her current peek lasted (Fig. 2b). This was true for the conditions where ants searched around a given location such as the nest or when released in completely unfamiliar location. In these conditions (BeN, ByN, U), the elapsed time, which necessarily correlates with the number of previous peeks, does not correlate with specific locations in the world (contrary to the ‘F’ condition). This shows that ants somehow gather information across time: either information about the overall time spent backwards or information about the number of peek previously displayed (see also Fig. 3c).

Former experiments have already shown that the behaviour of ants can be modulated by recent experience in the order of seconds to minutes – a form of hysteresis (Graham & Mangan, 2015). For instance, ants display a so-called backtracking behaviour only if they have recently perceived the nest surrounding (Wystrach, Schwarz, Baniel, & Cheng, 2013). Also, homing ants display higher meander in their paths when recapitulating a well-known route for the second time in a row (Collett, 2014; Wystrach, Schwarz, Graham, & Cheng, 2019). Finally, ants can remember the compass direction of a wind gust after being blown (Wystrach & Schwarz, 2013). In the present case, ants seem to also build up information about the recent past. Even though it makes functional sense it remains to be seen what neural or physiological mechanisms underlie this phenomenon.

### Conclusion

This study confirmed that ants walking backward are not just paying attention to celestial cues but combine multiple information from their PI, the recognition of terrestrial cues and temporal information such as the time they spent backward. All this information seems to be integrated in an overall directional drive. This drive, which reflects the current directional certainty, seems to (1) guide the ants backward path, (2) triggers peeking behaviour and (3) finally dictates whether or not to return to their food item during peeks. Importantly, this study shows that ants can recognise familiar terrestrial cues backward. In addition to the attractive memories facing toward the nest, the hypothesis that homing ants use a collection of repulsive visual memories facing away from the nest and possibly stored during their outbound trip was put forward. In the light of this hypothesis, visual navigation forward or backward can then simply be achieved by using the relative familiarity between both sets of opposite valence memories. As often with research on insect navigation, remarkably flexible behaviours incite researchers to endorse the simplest explanations (Wystrach & Graham, 2012).

## ACKNOWLEDGEMENTS

We are grateful for Xim Cerda and his helpful team at CSIC Seville for permanent assistance in logistics and administration during field work. We also thank Cornelia Buehlmann, Scarlett Dell-Cronin, Cody Freas and Michael Mangan for manual and moral support during field preparation and data collection. Finally, we are grateful for the constructive comments of Paul Graham on the manuscript. The study was partly financed by The European Research Council, 759817-EMERG-ANT ERC-2017-STG.

## AUTHOR CONTRIBUTIONS

SS, Conceptualization, Methodology, Validation, Formal analysis, Investigation, Writing – original draft preparation, Writing – review & editing, Visualization, Supervision, Project administration; LC, Investigation, Validation; EG, Investigation, Validation; AW, Conceptualization, Methodology, Validation, Formal analysis, Investigation, Writing – original draft preparation, Writing – review & editing, Visualization, Supervision, Project administration, Funding acquisition.

